# Identification of three distinct cell populations for urate excretion in human kidney

**DOI:** 10.1101/2023.06.29.545826

**Authors:** Yoshihiko M. Sakaguchi, Pattama Wiriyasermkul, Masaya Matsubayashi, Masaki Miyasaka, Nau Sakaguchi, Yoshiki Sahara, Minoru Takasato, Kaoru Kinugawa, Kazuma Sugie, Masahiro Eriguchi, Kazuhiko Tsuruya, Hiroki Kuniyasu, Shushi Nagamori, Eiichiro Mori

**Author notes:** Y.M.S., P.W. and M.M. contributed equally to this work. **Correspondence:** Shushi Nagamori. **Correspondence:** Eiichiro Mori.

## Abstract

In humans, uric acid is an end-product of purine metabolism. Urate excretion from human kidney is tightly regulated by reabsorption and secretion. At least eleven genes have been identified as human renal urate transporters. However, it remains unclear whether all renal tubular cells express the same set of urate transporters. Here we show that renal tubular cells are divided into three distinct cell populations for urate handling. Analysis of healthy human kidneys at single-cell resolution revealed that not all renal tubular cells expressed the same set of urate transporters. Only 32% of renal tubular cells were related to both reabsorption and secretion, while the remaining renal tubular cells were related to either reabsorption or secretion, at 5% and 63% respectively. These results provide physiological insight into the molecular function of the transporters and renal urate handling on cell-units. Our findings also suggest that three different tubular cell populations cooperate to regulate urate excretion from human kidney.

**Highlight/Key points:**

- We identified three distinct cell populations within the human renal anatomy that predict putative cellular transport mode, and our findings indicate cellular inhomogeneity with distinct roles such as urate secretion and reabsorption.
- Our model of physiological urate handling demonstrates the excretion dynamics in human kidney in terms of single cell-units.
- Our cellular urate transport analyses suggest the reversibility of some urate transporters even in certain physiological conditions.
- The physiological function of SLC2A9 is not limited to urate reabsorption; it is also involved in urate secretion restriction.
- This methodology can be applied to investigations of transport mechanisms in general, regardless of epithelial cell types, species, and substrates.

## Introduction

Excretion, the process of biological waste removal, is a vital homeostatic mechanism in all organisms. Humans have turnovers of 3.3 × 10^11^ cells per day (Sender & Milo, 2021), and wastes of nucleobases, the basic components of DNA and RNA, are removed with each turnover. Nucleobases are classified into pyrimidine and purine bases. Pyrimidine bases are catabolized to water, carbon dioxide, and ammonia, and purine bases are catabolized to uric acid as a final metabolic waste. Physiologically, more than 90% of serum uric acid exists as monosodium urate, which is excreted by urate transporters to maintain nucleic acid homeostasis. In other mammals, uric acid is hydrolyzed by uricase (urate oxidase). However, the loss of uricase during primate evolution (Kratzer *et al*., 2014) resulted in urate excretion becoming a more important nucleic acid homeostatic mechanism in humans than in other mammals.

Approximately two-thirds of urate excretion in humans is renal excretion, and the remaining one-third is extra-renal excretion, such as intestinal excretion (Sorensen, 1965). The renal excretion in humans is a reabsorption-dominant system composed of reabsorption and secretion (Steele & Rieselbach, 1967). Identification of urate post-secretory reabsorption site (Diamond & Paolino, 1973) led to these physiological urate excretion dynamics being proposed as a “four-component model” (Levinson & Sorensen, 1980). These dynamics are tightly regulated by functional urate transport unit (*urate transportome*), which consists of urate transporters and scaffold proteins (Anzai *et al*., 2004; Srivastava *et al*., 2019).

At least eleven genes have been identified as transporters in human kidney. The ATP-binding cassette (ABC) transporter or solute carrier (SLC) transporter (Enomoto *et al*., 2002; Ichida *et al*., 2003; Matsuo *et al*., 2008; Iharada *et al*., 2010; Chiba *et al*., 2015; Hyndman *et al*., 2016), and specific counterpart efflux transporters to influx transporters are involved in urate reabsorption and secretion, respectively (Bobulescu & Moe, 2012; Hyndman *et al*., 2016; Nigam & Bhatnagar, 2018). Recently, some urate reabsorption transporters (SLC22A11, SLC22A12, and SLC22A13) are found not to be co-expressed (Matsubayashi *et al*., 2021). However, it is still unclear whether all renal tubular cells express the same set of urate transporters, and how the distribution of their cells in human kidney result in physiological urate excretion.

To clarify whether all renal tubular cells express the same set of urate transporters, we analyzed healthy human kidneys at single-cell resolution. We found that not all the renal tubular cells expressed the same set of urate transporters, and identified three distinct cell populations for urate handling. Our findings revealed that the molecular function of the transporters is variable and differs across the expression patterns of *transportome* components on single cell-units. Based on these results, we developed a physiological model for urate handling on single cell-units to visualize the dynamics of urate excretion in human kidney with higher resolution.

## Methods

### snRNA-seq datasets

We made no distinction between single-nucleus RNA sequencing (snRNA-seq) or scRNA-seq and treat them as scRNA-seq datasets (Matsubayashi *et al*., 2021). We sought for scRNA-seq datasets based on two criteria. First, the datasets must be from kidneys of adult human males. Secondly, each dataset was used as a control in previous studies. The kidney in GSE118184 was validated as healthy kidney with a serum creatinine measurement of 1.03 mg/dL (Wu *et al*., 2018). Two kidneys in GSE131882 were validated as healthy kidneys via H&E images in which no evidence of glomerulosclerosis, interstitial fibrosis, or immune cell infiltrate was found (Wilson *et al*., 2019). Consequently, we selected and downloaded these three datasets from a public functional genomics data repository (Supplementary Table 1). As shown in other studies, we confirmed that we had an appropriate number of datasets for our study (Wilson *et al*., 2019; Liao *et al*., 2020). The gene name in the dataset “GSE118184” and “GSE131882” used HGNC gene symbol and Ensemble gene ID, respectively. We unified the gene names in all datasets as HGNC gene symbols by biomaRt (Durinck *et al*., 2005, 2009).

### Quality control of the datasets

Quality control was processed using the Seurat plug-in of R software (Butler *et al*., 2018; Stuart *et al*., 2019). Workflow was built according to the tutorial, “Guided tutorial – 2,700 PBMCs” listed in the Seurat website (https://satijalab.org/seurat/vignettes.html). To exclude low-quality and dying cells, we filtered cells out if their mitochondrial gene content was > 5% because high mitochondrial gene content is reported to be related to low-quality or dying cells. To exclude empty droplets and cell doublets or multiplets, we filtered cells out if their gene counts detected were less than 200 or more than each threshold. The thresholds of the three datasets were set as 6000, 6000, and 3000, respectively. Finally, the datasets used in this study consist of three males and a total of 10,080 cells (Supplementary Table 1).

### Multiple dataset integration and data pre-processing of cell clustering

Data integration and data pre-processing of cell clustering were performed using the Seurat plug-in of R software. The workflow was set following the tutorials, “SCTransform”, “Cell Cycle Regression”, and “Integration and Label Transfer - SCTransform” listed in the Seurat website (https://satijalab.org/seurat/vignettes.html). We selected sctransform as a normalization method; it uses regularized negative binomial regression to normalize UMI count data (Hafemeister & Satija, 2019). Variations in technical factors of scRNA-seq data tend to confound the actual biological variations (Stegle *et al*., 2015; Vallejos *et al*., 2017). We utilized the sctransform normalization and data integration by Seurat in the pre-processing stage to reduce more technical factor variations and preserve real biological variations compared to the standard Seurat workflow (Butler *et al*., 2018; Hafemeister & Satija, 2019). We performed the sctransform normalization twice. The first sctransform normalization was performed with the regression of nFeature_RNA and nCount_RNA.

Cell cycle regression is a method to lessen the effects of cell cycle heterogeneity in scRNA-seq data by calculating cell cycle phase scores based on canonical markers (Nestorowa *et al*., 2016). The calculation utilized the values obtained after the first sctransform. After the calculation of cell cycle scores, a second sctransform normalization was performed again with the regression of nFeature_RNA, nCount_RNA, and cell cycle scores. Finally, we integrated the three datasets according to the tutorial “Integration and Label Transfer - SCTransform” (Butler *et al*., 2018).

### Cell clustering

To visualize the cell types in the three datasets, we performed cell clustering. The process of cell clustering was also performed using the Seurat plug-in of R software. The workflow was built according to the tutorial “Integration and Label Transfer - SCTransform” in the Seurat website (https://satijalab.org/seurat/vignettes.html). After an integration of the three datasets, we projected these datasets onto two dimensions with uniform manifold approximation and projection (UMAP), a dimension reduction technique. All single nuclei were divided into a total of 16 clusters. The clusters were named by the renal anatomical, which were annotated by marker genes (Supplementary Table 2). The marker genes were selected based on criteria listed in previous studies (Wu *et al*., 2018; Wilson *et al*., 2019).

### Annotation of segments in proximal tubule (PT) clusters

Three PT clusters were divided into S1: S1 segment of proximal tubule, S2: S2 segment of proximal tubule, and S3: S3 segment of proximal tubule. These clusters were annotated by marker genes previously reported (Supplementary Table 3). Selection of the marker genes for S1 and S3 segments were based on previous studies (Ba *et al*., 2003; Mather & Pollock, 2011). As no specific marker genes for S2 segments were reported, we annotated the S2 segments from the cluster which was not marked as S1 or S3 segments in three PT clusters.

### Gene positivity and gene negativity

Gene expression data were normalized by the “LogNormalize” method on Seurat. First, gene counts for each cell were divided by the total counts for that cell and the values multiplied by 10,000 were set as the default scale factor on Seurat. Then, these were natural-log transformed. After the log transformation, we defined “gene X”-positivity (the presence of gene X) as cells that expressed “gene X” above the cutoff value. By contrast, “gene X”-negativity (the absence of gene X) was defined as cells that expressed “gene X” less than the cutoff value. A value ‘0.5’ was set as the cutoff value because the expression levels had a neckline around the value shown in a violin plot. The gene-positive cell numbers were counted after the translation from the quantitative expression data to binary expression data (gene-positive or negative).

All urate transporter genes were classified into one of four types, Apical Influx (AI) transporter, Apical Efflux (AE) transporter, Basolateral Influx (BI) transporter, or Basolateral Efflux (BE) transporter, based on a previous study (Hyndman *et al*., 2016). The positivity or negativity of each gene was then examined as described above. After that, their expression profiles were constructed into cell populations based on the positivity/negativity of AI and BI transporters. We classified cell populations into one of four cell populations (Apical Influx Transporter (AIP) cell population, Basolateral Influx Transporter (BIP) cell population, Double Positive (DIP) cell population, or Double Negative (DIN) cell population). The AIP cell population had the positivity of at least one AI transporter (*SLC22A11* and/or *SLC22A12*) and the negativity of BI transporters (*SLC22A6*, *SLC22A7*, and *SLC22A8*). The BIP cell population had the positivity of at least one BI transporter (*SLC22A6*, *SLC22A7*, and/or *SLC22A8*) and the negativity of AI transporters (*SLC22A11* and *SLC22A12*). The DIP cell population had the positivity of both AI transporters (*SLC22A11* and/or *SLC22A12*) and BI transporters (*SLC22A6*, *SLC22A7,* and/or *SLC22A8*). The DIN cell population had the negativity of both AI and BI transporters. After that, we subcategorized each cell populations by adding the positivity/negativity of AE and BE transporters in the same way. Finally, we calculated the cell number presented in Table 2 as the numbers of cells which showed the positivity of indicated transporters.

### Visualization of the expression data – dot plots, violin plot, and bar plot

The visualization of the gene expression data was performed using Seurat plug-in of R software (Butler *et al*., 2018; Stuart *et al*., 2019), ggplot2 plug-in of R software (Wickham, 2010), and matplotlib.plt library of python software (Hunter, 2007). All expression levels used in data visualization were the scaled data after the log transformation as described in the previous section. Dot and violin plots of gene expression in each cluster and scatter plots of two genes expression were created by Seurat function, *“Dotplot”*, *“VlnPlot”*, plug-in of R software. We created bar plots using matplotlib, which is the Python library for producing plots and data visualizations. Plots reside within a *Figure* object that was created by a function named *“Figure”* in matplotlib.plt library, and the bar plots were made by the *show* method in *Figure* object.

## Data availability

R and Python codes for data analysis and visualization is available in GitHub repository at https://github.com/yoshi-sci/urate-transporter-analysis.

## Results

### 1 Urate handling transporter identification and localization

To examine the localization of the transporters in human kidney at single-cell resolution, we first verified the renal anatomical regions of all single nuclei in the three datasets. After unsupervised clustering of all single nuclei in the datasets, all clusters were annotated by the marker genes (Figure 1A, Supplementary Figure 1A). The annotation was based on the anatomical regions, and cells in the clusters were calculated as percentages in the regions (Figure 1B, Supplementary Figure 1B). Proximal tubule (PT) is the region for urate reabsorption and secretion, and it was divided into three segments based on anatomical localization. Using the PT region-specific marker genes, we were able to annotate three PT segments: S1, S2, and S3 segments (Figure 1C). The three analyzed datasets covered all cell types of renal tubules including three PT segments (Supplementary Figure 1C), which allowed us to evaluate the localization of urate transporters.

**Figure 1.**
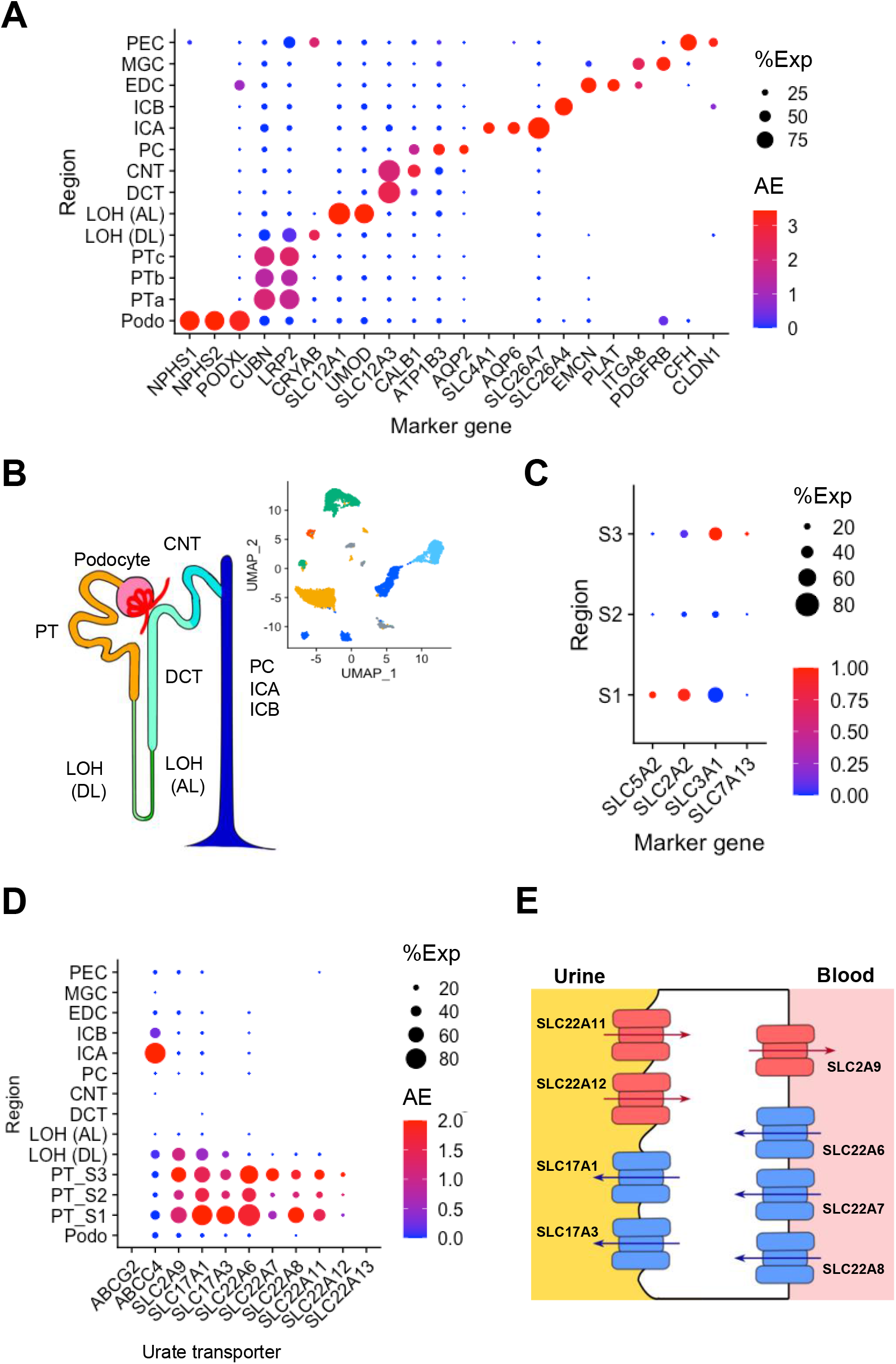
Identification of regions and transporters for urate handling in human kidney. (A) Annotation of the 14 clusters by the marker genes. Podo, podocyte; PTa-c, three clusters of proximal tubule; LOH (DL), the loop of Henle (descending loop); LOH (AL), the loop of Henle (ascending loop); DCT, distal convoluted tubule; CNT, connecting tubule; PC, principal cell; ICA, intercalated cell type A; ICB, intercalated cell type B; EDC, endothelial cell; MGC, mesangial cell; and PEC, parietal epithelial cell. (B) A schematic diagram of human nephron anatomy and unsupervised clustering of healthy human adult renal cells. (C) The dot-plot of PT marker genes: S1 markers and S3 markers. S1-S3, S1-S3 segments of proximal tubules. Dot sizes indicate the gene frequencies (%Exp) and dot colors indicate average expression levels (AE) in the cluster. (D) The dot-plot of the gene expression of urate transporters in the renal regions. (E) A schematic diagram of urate transporters expressed in the PT and DL clusters. The apical membrane is to the left of the cells and the basolateral membrane is to the right. Red symbols represent transporters which are reconstituted to urate reabsorption. Blue symbols represent transporters which are reconstituted to urate secretion. Arrows indicate directions of urate flow.

At least eleven genes have been identified as urate transporters in human kidney. To confirm the transporters and the localization for urate handling in human kidneys, we focused on the expression of the eleven known transporters across the regions (Supplementary Figure 1D). Except for *ABCG2*, *ABCC4,* and *SLC22A13*, the remaining eight transporters were specifically expressed in the PT and the descending loop of Henle (DL). *ABCG2* and *SLC22A13* were rarely expressed in all clusters, while *ABCC4* was the only transporter highly expressed in the intercalated cell type A region (Figure 1D). The results clarified that transport via these eight transporters existed mainly in the PT, and to some degree in the DL region. Consequently, our subsequent analyses focused on 1) the four regions (S1, S2, S3, and DL), and 2) the eight urate transporters specifically expressed in the clusters (SLC2A9, SLC17A1, SLC17A3, SLC22A6, SLC22A7, SLC22A8, SLC22A11, and SLC22A12) (Figure 1E).

The eight transporters have unique characteristics for cellular urate transport. In physiological conditions, the transporters are proposed to be unidirectional based on the transport affinity and substrate availability both outside and inside the cells (Bobulescu & Moe, 2012; Hyndman *et al*., 2016). Table 1 summarizes the eight transporters in terms of their transport systems, substrates, affinities (K_m_) for urate, and average expressions in the clusters of PT and DL (Koepsell, 2013; Mueckler & Thorens, 2013; Reimer, 2013), demonstrating that these transporters can be classified based on their cellular function for urate transport.

**Table 1.**
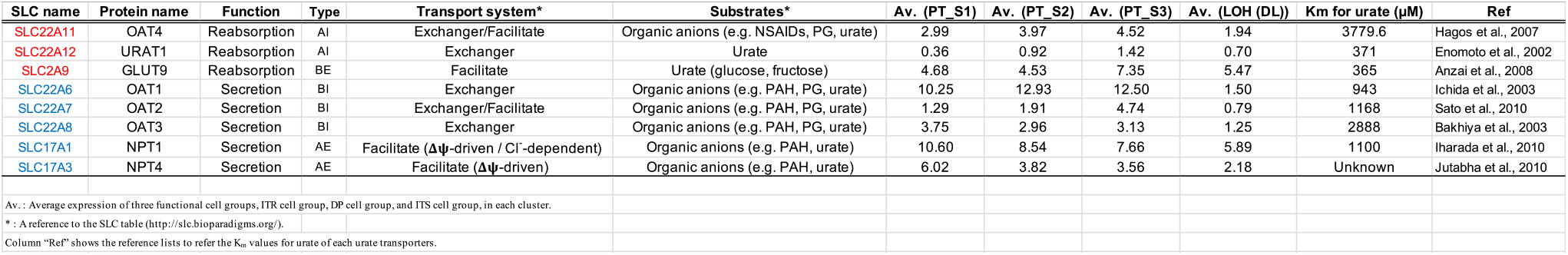
Transport properties of urate transporters expressed in renal tubular regions of human males in this analysis.

### 2 Classification of four transporter types and four cell populations

To classify the eight transporters, we considered polarized localization (apical or basolateral membranes) and urate flux (efflux or influx). We categorized the eight transporters into one of four types (Figure 2A): Apical Influx urate transporters (AI transporters), Apical Efflux urate transporters (AE transporters), Basolateral Influx urate transporters (BI transporters), and Basolateral Efflux urate transporters (BE transporters). The cooperation of AI and BE transporters leads renal tubular cells to function as reabsorption, whereas the cooperation of BI and AE transporters leads their cells to function as secretion. This method of classification helped us to predict the cellular urate transport function.

**Figure 2.**
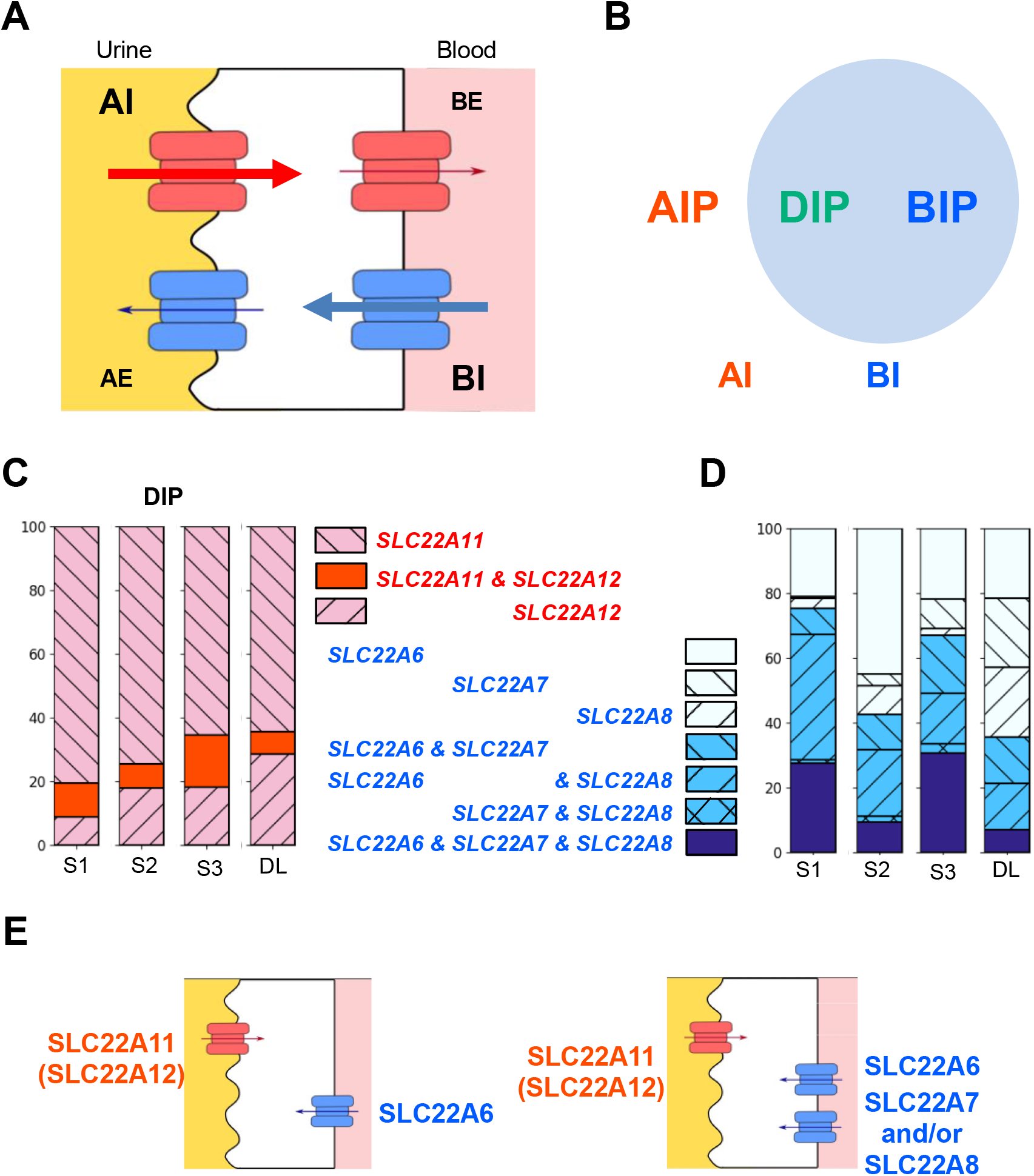
Classification of transporter types and cell populations for urate handling. (A) A schematic diagram of four types of urate transporters classified based on their subcellular localization and physiological roles (apical/basolateral and influx/efflux). The eight transporters were classified into one of the following four transporter types: (1) SLC22A11 and SLC22A12 as AI transporters; (2) SLC17A1 and SLC17A3 as AE transporters; (3) SLC22A6, SLC22A7, and SLC22A8 as BI transporters; or (4) SLC2A9 as BE transporter. The cooperation of an Apical Influx (AI) transporter and a Basolateral Efflux (BE) transporter leads to urate reabsorption (red). The cooperation of a Basolateral Influx (BI) transporter and an Apical Efflux (AE) transporter leads to urate secretion (blue). (B) Venn diagram indicating cell populations, which were classified based on the positivity of influx transporters. Cell clusters in the PT and DL were classified into one of the following cell populations: AIP cell population, BIP cell population, DIP cell population, or DIN cell population. (C-D) Bar plots indicating the percentage (y-axis) of AI transporters (C) and BI transporters (D) in the DIP cell population across the regions (x-axis) in the DIP cell population. (E) Expression pattern schema of influx transporters. The apical membrane is to the left of the cells and the basolateral membrane is to the right of the cells. Red symbols represent the AI transporters, which are only single kind expressed in single cells. Blue symbols represent the BI transporters, which are single or multiple kind(s) expressed in single cells. Arrows indicate directions of urate flow.

Based on the cellular expression profiles of the transporter types, we hypothesized that cell populations have distinct cellular functions for transport. Based on the gene-positivity of the influx transporters, PT and DL cells were categorized into one of four cell populations (Figure 2B): 1) AI transporter—positive cell population expressing AI transporters but not BI transporters (AIP cell population); 2) BI transporter—positive cell populations expressing BI transporter but not AI transporters (BIP cell population); 3) Double Influx transporters—positive cell populations expressing both AI and BI transporters (DIP cell population); and 4) Double Influx transporters—negative cells lacking the expression of both AI and BI transporters (DIN cell population). For example, the AIP cell population expressed *SLC22A11*, *SLC22A12*, or both, but had no expression of *SLC22A6*, *SLC22A7,* or *SLC22A8*. The positivity of AI (BI) transporters was defined as the expression of at least one AI (BI) transporter in each cell.

To elucidate the expression profiles of the influx transporters in the cell populations, we first examined how frequently the same types of influx transporters were co-expressed. The most frequent AI transporter was *SLC22A11* across the regions (Supplementary Figure 2A, B), while the most frequent BI transporter was *SLC22A6* (Supplementary Figure 2C-E). We found that for AI transporters, all tubular regions showed that about 70 – 80% of the DIP cell population expressed *SLC22A11* but not *SLC22A12*, and less than 10% of the DIP cell population expressed both *SLC22A12* and *SLC22A11* (Figure 2C). For BI transporters, approximately 75% of the DIP cell population in the S1 and S3 regions showed co-expression of two or more different BI transporters, although the proportion decreased in the S2 and DL regions (Figure 2D). These frequency trends of influx transporters in the AIP (BIP) cell population were also found to be common in the DIP cell population (Supplementary Figure 2F, G). These results suggest that only one kind of AI transporter, mostly SLC22A11, is expressed, while other kinds of BI transporters might be co-expressed in addition to SLC22A6 (Figure 2E).

### 3 Expression of the efflux transporters indispensable for predicting cellular urate transport

We made the following two assumptions when classifying the three cellular urate transport directions, reabsorption mode, secretion mode, and bidirectional mode. First, we assume that transport direction was based on the availability of urate in both the lumen and the blood, and the efflux of urate would not take place without the influx of urate into the cells. Thus, the DIN cell population was omitted due to the lack of influx transporters (Supplementary Figure 3A). Secondly, we assumed that a complete transport direction mode required the co-expression of an influx transporter on one side of the polarized membrane (either AI or BI transporter) and an efflux transporter on the other side of the polarized membrane in the same cell. In other words, the AI and BI transporters were assigned as the primitive transporters, while the AE and BE transporters accomplished such transport modes as the derivative transporters. These two assumptions were applied to the determination of the cellular directional mode in the three cell populations: AIP, BIP, and DIP cell populations (Figure 3A).

**Figure 3.**
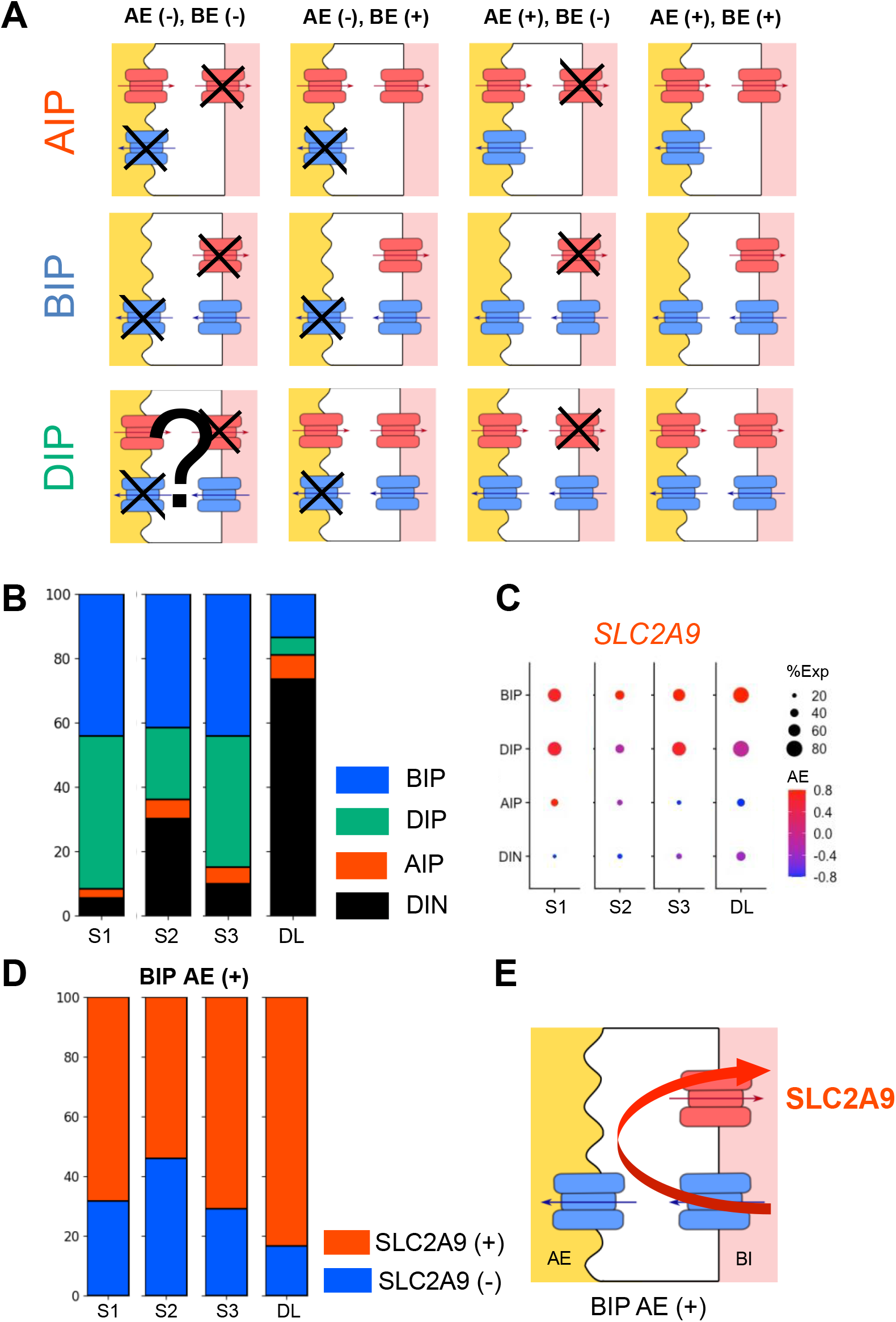
Prediction of cellular urate transport direction from the expression of the transporters. (A) Models indicating all potential expression patterns of urate transporters in each of the following cell populations: AIP cell population, BIP cell population, and DIP cell population. Colors indicate types of transporters: red, AI or BE transporters; blue, BI or AE transporters. Arrows show urate transport directions. (B) Proportion of the cell populations in the regions. The bar plot shows the percentage of cells (y-axis) in each cell population across the PT and LOH (DL) clusters (x-axis). (C) The dot-plot indicates the frequency and expression levels of SLC2A9 (x-axis) across the cell populations (y-axis) along the three PT segments (S1-S3) and DL. Dot sizes refer to the frequency of a molecule expressed in the cell population (%Exp), while dot colors indicate expression levels (AE). (D) Positivity proportion of SLC2A9 in the BIP cell population with the AE transporters. The bar plot shows the percentage of cells (y-axis) in each cell population across the PT and LOH (DL) clusters (x-axis). (E) A schematic diagram of SLC2A9 cell function in the BIP cell population. The apical membrane is to the left of the cells and the basolateral membrane is the right of the cells. Red symbols represent the AI transporters. Blue symbols represent the BI and AE transporters. Arrows indicate directions of urate flow.

To visualize urate handling across the regions, we clarified which cell populations were present in the regions. The BIP and DIP cell populations were highly dominant in the S1 region and gradually decreased along the anatomical region (Figure 3B). This suggests that the three cell populations are not only in the PT region but also in the DL region, and that their proportion decreases as anatomical localization progressed downward from the S1 to DL regions. In all PT segments, the proportion of the BIP cell population was approximately 50%, whereas the AIP cell population was the smallest among the cell populations (Figure 3B). These analyses indicate that the cells expressing AI transporters (the AIP and DIP cell populations) also express the BI transporters, but the cells expressing BI transporters (the BIP and DIP cell populations) do not necessarily express AI transporters.

To elucidate the expression profiles of the efflux transporters across the cell populations, we analyzed the frequencies and expression levels of BE and AE transporters (Figure 3C, Supplementary Figure 3B, C). In all PT and DL clusters, the BIP and DIP cell populations exhibited a high frequency and expression level of *SLC2A9*, the only known BE transporter, while the AIP cell population showed a low frequency and expression level of *SLC2A9* (Figure 3C). These results suggest that SLC2A9 is expressed not only in the renal tubular cells related to cellular urate reabsorption (the DIP and AIP cell populations), but also in the renal tubular cells involved in cellular urate secretion (the BIP cell population); SLC2A9 plays the role of lowering the secretion efficiency. To clarify the regions affected by SLC2A9 to decrease secretion efficiency, we showed *SLC2A9* expression in the BIP cell population with AE transporters. The expression had peaks on both ends, the S1 and DL regions (Figure 3D), suggesting that SLC2A9 lowers the secretion efficiency in the BIP cell population, especially at the beginning and end of urate handling regions (Figure 3E).

### 4 Contribution of urate transportome to urate handling

We considered the presence of regulatory factors that promoted the expression patterns of the transporters. We hypothesized that one of the key factors was PDZ proteins, which are scaffold proteins that interact with various transmembrane proteins, including transporters via PDZ motifs at the C-termini. PDZK1, a scaffold protein, forms the functional urate transport unit (*urate transportome*) by clustering urate transporters at the apical membrane (Anzai *et al*., 2004; Srivastava *et al*., 2019). We first investigated the correlation between PDZK1 expression patterns and the cell populations. The expression of *PDZK1* in each cell population revealed that the DIP cell population had the highest frequency and expression level of *PDZK1;* PDZK1-positive cells accounted for approximately 60% of the DIP cell population (Figure 4A), suggesting that the DIP cell population expresses PDZK1 more frequently and strongly than the other cell populations.

**Figure 4.**
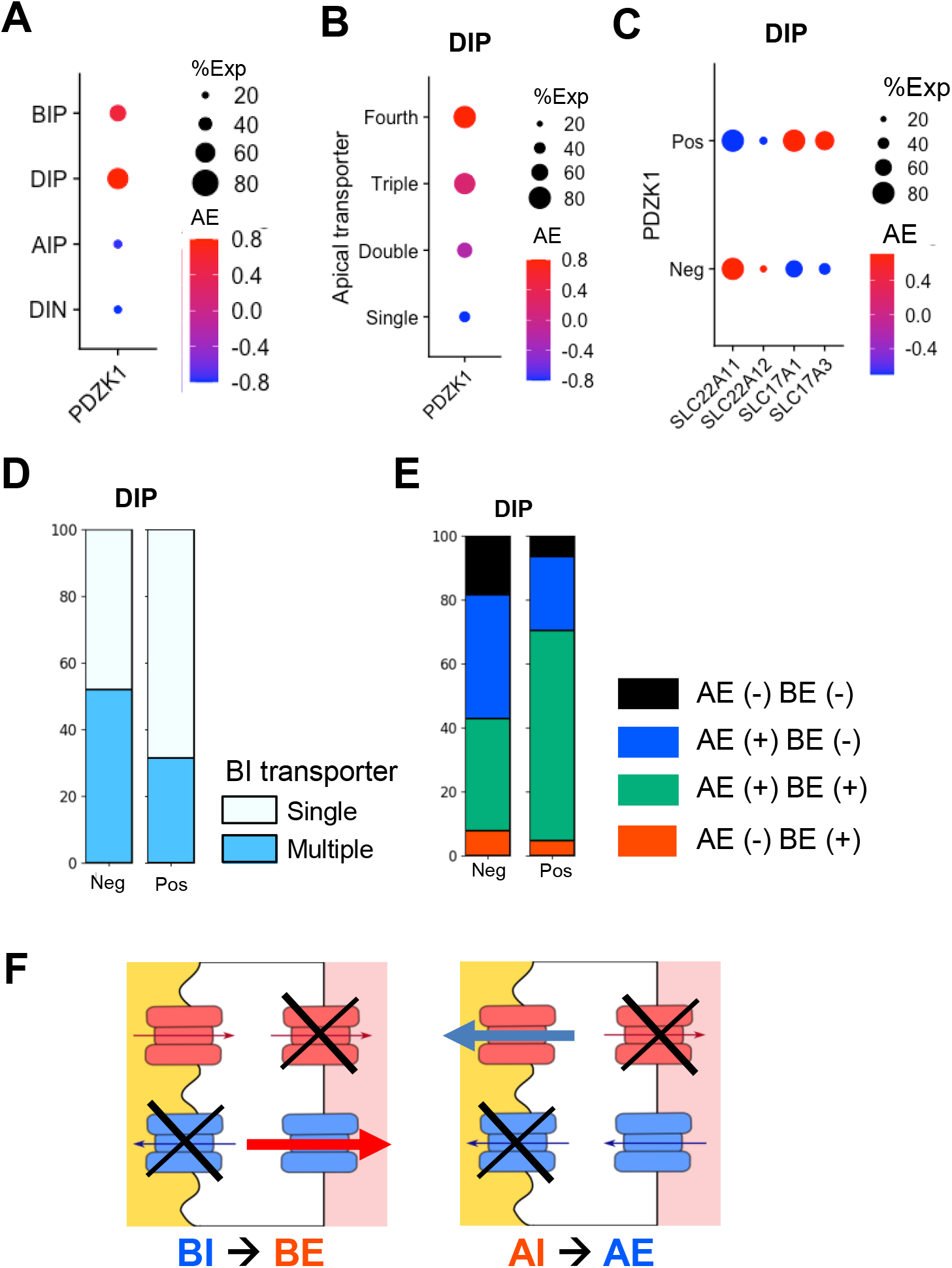
Expression profiles of urate transporters and PDZK1 in DIP cell population. (A) Expression levels and frequencies of P DZK1 in the cell populations. (B) Expression levels and frequencies of PDZK1 in the DIP cell population are rearranged by the numbers of apical membrane transporters. (C) The dot-plot indicates the frequency and expression levels of the apical transporters (x-axis) across the positivity of PDZK1 (y-axis). Dot sizes refer to the frequency of a molecule expressed in the cell population (%Exp), while dot colors indicate the expression levels (AE). (D) Cell proportion of the BI transporter kind numbers. The bar plot shows the percentage of cells (y-axis) across the positivity of PDZK1 (x-axis). (E) Cell proportion by the efflux transporters (AE and BE transporters). The bar plot shows the percentage of cells (y-axis) across the positivity of PDZK1 (x-axis). (F) A schematic diagram of transporter reversibility from influx transporter to efflux transporter in the DIP cell population. the apical membrane is to the left of the cells and the basolateral membrane is the right of the cells. Red symbols represent the AI and BE transporters. Blue symbols represent the BI and AE transporters. Arrows indicate directions of urate flow.

In contrast to the BIP and AIP cell populations, the DIP cell population potentially directs both secretion and reabsorption and changes the net cellular transport mode by the transporters expressed. To clarify whether the PDZK1-mediated urate *transportome* favors the expression patterns of the transporters, we showed the expression of *PDZK1* over the number of apical transporters. The more diverse the apical urate transporters were, the higher the expression levels and frequencies of *PDZK1* were (Figure 4B). *PDZK1*-positive cells had higher or equal frequencies of all apical transporters compared to *PDZK1*-negative cells, and the AE transporters (*SLC17A1* and *SLC17A3*) were also more strongly expressed in *PDZK1*-positive cells (Figure 4C). These results indicate the contribution of PDZK1 to the formation of the cellular urate *transportome* at the apical membranes of the DIP cell population.

To demonstrate that the positivity of PDZK1 is related to the net cellular transport mode in the DIP cell population, we focused on the expression of the basolateral transporters in addition to the apical transporters across the positivity of *PDZK1*. As shown in Figure 2E, the DIP cell population had single or multiple kinds of BI transporters. *PDZK1*-positive cells more frequently had multiple kinds of BI transporters than *PDZK1*-negative cells (Figure 4D). Also, *SLC2A9,* the AE transporter, was expressed more frequently and strongly in *PDZK1*-positive cells (Supplementary Figure 4A). About 60% of the *PDZK1*-positive DIP cell population accounted for the positive expressions of both efflux transporter types (the AE and BE transporters), while the *PDZK1*-negative DIP cell population more frequently accounted for the negative expressions of the efflux transporter types, including the influx transporter-only population (Figure 4E). These results suggest that PDZK1 has a positive relationship with the basolateral transporters in addition to apical transporters and changes the net cellular transport mode in the DIP cell population.

According to our cell population classification criteria, efflux transporters are not always expressed in the DIP cell population. In fact, about 20% of the PDZK1-negative DIP cell population expressed none of the efflux transporters (Figure 4E). It is interesting to consider the kind of cellular transport modes the DIP cell population had without efflux transporters. Even if the cell population influxes urate at an early stage, it would generate higher and higher intracellular urate concentration. We expect that the change in intracellular and extracellular concentration gradient would then subsequently drive the role of the influx transporters to change the efflux physiologically. The fundamental principle of transporters is that the transport direction is determined by the substrate concentration gradient. The DIP cell population without efflux transporters could potentially switch the direction of urate transport from influx to efflux, for example, from a BI transporter to a BE transporter or from an AI transporter to an AE transporter (Figure 4F).

### 5 Design of a triple cell-unit model of urate handling, the “*CUTE model*”

The *Cellular Urate Transport Excretion* (*CUTE*) *model* proposed three representative cell populations: the BIP, DIP, and AIP cell populations (Figure 5A). The cellular transport modes were defined by the cell populations (either secretion or reabsorption); if the BIP (or AIP) cell population expressed AE (BE) transporters, the population worked for urate secretion (reabsorption) (Figure 5A). The DIP cell populations could work for urate secretion or reabsorption depending on the transporter types expressed. The DIP cell population with either AE (or BE) transporters could potentially play a role in urate secretion (or reabsorption), while the other DIP populations (“AE-BE-” or “AE+ BE+”) could work bi-directionally (Supplementary Figure 5A). Our results suggest that not all renal tubular cells express the same set of urate transporters, and that the cells involved in the cellular reabsorption are distinct from those involved in the cellular secretion.

**Figure 5.**
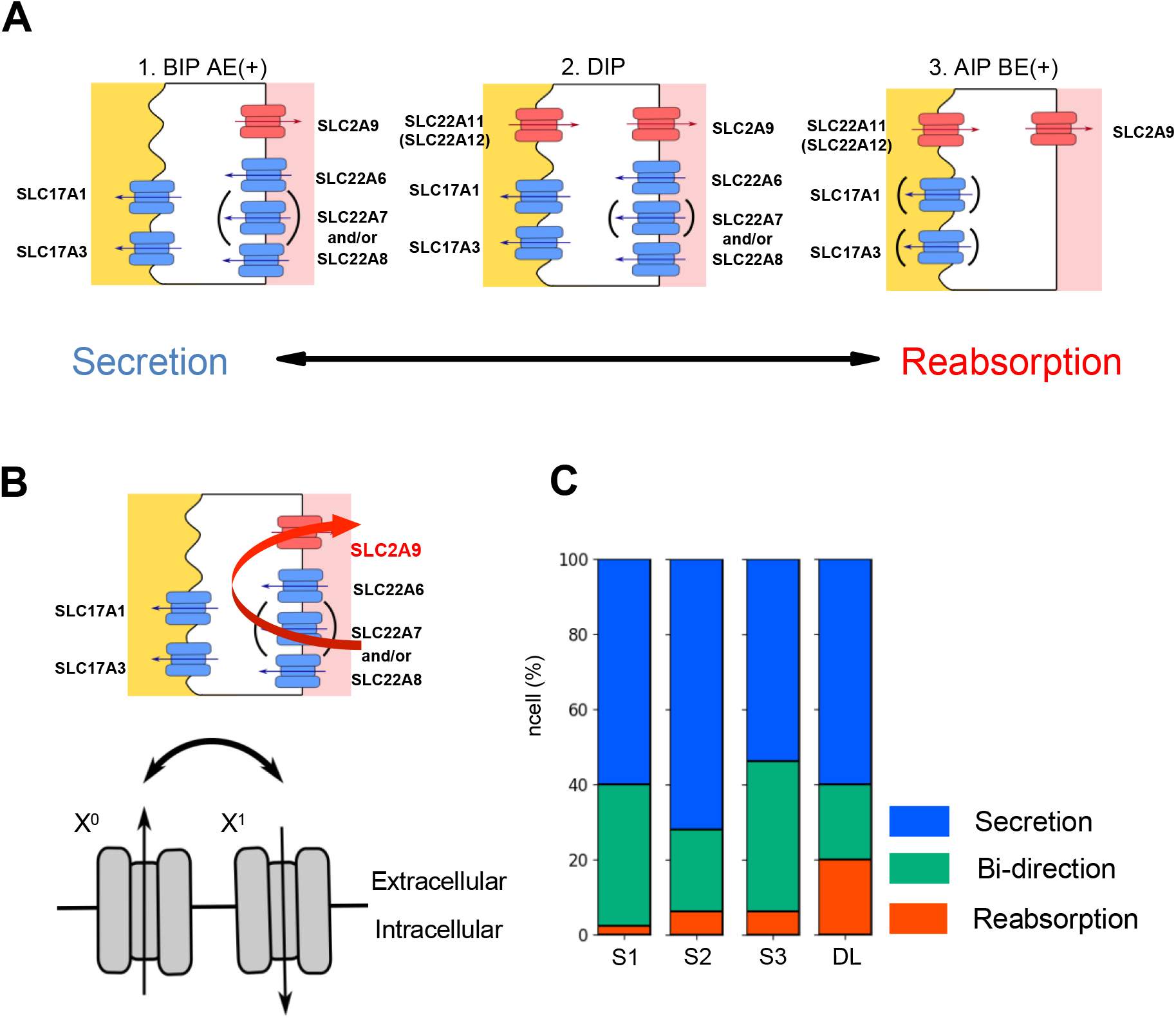
Major cell populations explaining renal urate transport dynamics. (A) Representative cell populations that contributed to urate reabsorption and secretion. Colors indicate types of transporters: red, AI or BE transporters; blue, BI or AE transporters. (B) A schematic diagram illustrating the feasibility of the superiority of urate reabsorption over urate secretion. Upper panel: Secretion attenuation of BIP cell population by SLC2A9 at all tubular regions. Lower panel: Reversibility of the transporter function between urate influx and efflux in human kidney. (C) The bar plot showing cell proportion by cellular urate handling in three PT segments and LOH (DL) regions.

The clarification of the set of urate transporters at single-cell resolution provided some insights into transporter functions. First, the role of SLC2A9 could be significant to lower the secretion efficiency, which supports the clinical data of low urate excretion (Figure 5B: Upper panel). Second, the *CUTE model* proposed a property of transport reversibility in some transporters in physiological conditions (Figure 5B: Lower panel). These results were found only when the molecular function was taken into consideration on cell-units.

To clarify the cell proportion for urate handling across the regions, we quantified the counts for reabsorption cells, bi-directional cells, and secretion cells (Figure 5C, Supplementary Table 4). Proportions of the regions were shown (Supplementary Figure 5B), and the average proportions of the cell populations in the regions were as follows: secretion 63%, bi-direction 32%, and reabsorption 5%. The ratio of reabsorption cells per secretion cells had an upward trend along the renal tubule (S1 0.04, S2 0.09, S3 0.12, and DL 0.33); in particular, the ratio in the DL region was by far the highest among the regions. These results suggest that urate reabsorption is not as superior as urate secretion at the beginning of PT but becomes more significant at S3 and DL. Accordingly, secretion could be most active at the beginning of the PT and gradually declines along the PT segments.

## Discussion

The concept of having same sets of multiple transporter molecules in the same cells has been proposed (Bobulescu & Moe, 2012; Hyndman *et al*., 2016). Our study elucidated these sets of the transporters in single cells and predicted the directions of the cellular urate transport. Four factors were used to construct the cellular model. The first factor was the identification of renal anatomical regions where known urate transporters were expressed, and the regions were expected to be the functional regions for the reabsorption and secretion. The next factor was the identification of four transporter types, which were defined based on the characteristics of the transporters. The third factor was the classification of four cell populations based on the expression profiles of the transporters at single-cell levels. The combination of the transporter types and the cell populations led to the prediction of cellular direction modes of urate transport as the fourth factor. The *CUTE model* confirmed the expression of the transporter molecules, except SLC22A13, ABCG2, and ABCC4, in human kidneys. In addition, the *CUTE model* clarified the cellular inhomogeneity (Figure 5A) and complexity of cell population distribution along the anatomical regions (Figure 5C). An advantage of having inhomogeneous cell populations is probably the modulation of urate homeostasis.

The *CUTE model* proposes some concepts about the molecular function of transporters. One is a reversal property of some transporters in human kidneys (Figure 5B: Lower panel). Bi-directional functions of urate transporters *in vitro* have been previously reported (Enomoto *et al*., 2002; Bakhiya *et al*., 2003; Anzai *et al*., 2008; Jutabha *et al*., 2010) while the urate transporters in physiological conditions are proposed to restrict in uni-directional transport based on the transport affinity and substrate availability at both the outside and inside of the cells (Bobulescu & Moe, 2012; Hyndman *et al*., 2016). The *CUTE model* suggests that the urate transporters function bi-directionally *in vivo* as well as *in vitro*.

The reversibility is not applied to all urate transporters given in previous reports. For example, SLC22A7 and SLC22A11 were reported to be unidirectional urate transporters (Hagos *et al*., 2007; Sato *et al*., 2010). In addition, SLC22A12 and SLC2A9 tend to cooperate in urate reabsorption in the co-expressing cells (Nakanishi *et al*., 2013), although both transporters possibly divert in opposite directions (Enomoto *et al*., 2002; Anzai *et al*., 2008). Cumulatively, these reports suggest that BI and AE transporters, except SLC22A7, are capable of switching their transport directions, whereas AI and BE transporters have restricted transport directions. The *CUTE model* relatively restricts the direction modes yet accommodates some flexibility toward the reabsorption mode. Urate reabsorption is superior to urate secretion in the whole kidney. This model suggests a mechanism to keep the superiority of urate reabsorption.

Knowledge from human samples is essential to truly understanding human physiological systems. The *CUTE model* not only explains the molecular significance of urate transporters but also proposes clinical relevance. Currently, only two urate transporters related to the Mendelian disorders have been reported: SLC22A12 (OMIM 220150) which is responsible for renal hypouricemia type 1 (RHUC1) (Sperling, 2006), and SLC2A9 (OMIM612076) which is responsible for renal hypouricemia type 2 (RHUC2) (Kawamura *et al*., 2011; Chiba *et al*., 2014). Before the characterization of SLC2A9, most of the clinical studies focused on SLC22A12, and over 90% of renal hypouricemia was reported from *SLC22A12* mutations (Ichida *et al*., 2004). Here, we refer to the clinical association of SLC2A9, especially in the elderly male population, because the datasets used in this study were from elderly males. The *CUTE model* advocates SLC2A9 over other transporters because of its single-player BE transporter, high expression profiles, and dual functions in reabsorption and secretion attenuation (Figure 5B: Upper panel). Future analysis of more datasets covering a wide range of ages will provide additional clues about other urate transporters.

The *CUTE model* provides some clues as to where novel urate transporter candidates might be located. Our model suggests the existence of novel transporter(s) involved in urate reabsorption. The most possible candidate is a novel BE transporter, which is likely expressed more frequently and strongly in the AIP and DIP cell populations than in the BIP and DIN cell populations due to the weak expression of SLC2A9 in current AIP and DIP cell populations within the S2 region (Table 2). The model itself can provide a ready platform for further analysis when a novel urate transporter is identified. The analytical model from this study is beneficial for both before and after the discovery of candidate transporters.

The series of analyses presented here is highly scalable, and the methodology itself can be a prototype for other metabolite analyses in all organs. It is well known that multiple transporters in the same cells coordinate to handle a substance across a tissue region, such as nephrons, intestinal epithelia, liver tissues, and blood-brain barrier. Our analytical strategy is useful for broad types of transporters and substrates, leading to the understanding of molecular transport mechanisms in actual physiological systems. Our study suggests the functional cooperation of SLC22A11 and SLC22A6 for urate reabsorption (Figure 5A). This functional cooperation is also the canonical transport pathway for other endogenous anion substrates or drugs (Hagos *et al*., 2008, 2012; Willemin *et al*., 2021). The methodology employed in this study is potentially a strong tool to elucidate the transport mechanism in physiological and pathological systems.

In interpreting the results, it is important to understand the limitations of the scRNA-seq data we used. We did not consider factors such as race/ethnicity, sex, and age, although they have been reported to impact serum uric acid levels (Barry *et al*., 1992; Julius *et al*., 2004). These three factors may cause different expression levels and/or frequencies of urate transporters, as seen in the case of SLC22A12 (Joseph *et al*., 2015). Recently, SLC22A13 was found to contribute to urate reabsorption in an elderly Japanese male (Toyoda *et al*., 2022), but the non-Asian scRNA-seq data we used in this study suggests that the low contribution of SLC22A13 is due to its low expression frequency (Figure 1A, Supplementary Figure 2B). In addition, as sequencing techniques and analyses are further developed, metadata analyses including these factors are expected. Further study is required to address these issues.

In conclusion, the *CUTE model* showed three representative cell populations with distinct transporter expression patterns and the differential distribution of the cell populations along the tubular regions. The construction of the model clarified that the molecule function of the transporters was variable and differed across the expression patterns of the transporters on single cell-units. The component of urate excretion was from 1) reabsorption cells (5% of the cell population) which certainly drive urate reabsorption; 2) secretion cells (63%) which are supposed to drive urate secretion, but whose process could be attenuated by the high affinity of SLC2A9; and 3) bi-directional cells (32%) which potentially function as both reabsorption and secretion, but tentatively drive the reabsorption over the secretion by the reversibility of some transporters. The inhomogeneity of the cell populations allowed us to predict distinct net cellular transport modes, suggesting that the cells for urate reabsorption are relatively different from the cells for urate secretion. The distribution indicates the most active region for urate transport is at proximal tubules and to some degree at the descending loop of Henle, which lead us to predict urate excretion dynamics in certain regions of the nephrons. Overall, the urate transport dynamics described by our model indicate the superiority of urate reabsorption over urate secretion, leading to high uric acid excretion. Our methodology reflects the basic concept that any process, whether secretion or reabsorption, must pass through at least two types of transporters: one at the apical membrane and the other at the basolateral membrane. We believe our model can be applied to the analysis of transporter systems in general, bridging the gap from molecular function to cellular function.

## Author Contributions

**Y.M.S.**: Conceptualization, Formal analysis, Investigation, Software, Writing (original draft), Writing (review and editing). **P.W.**: Conceptualization, Funding acquisition, Writing (original draft), Writing (review and editing). **M.Ma.**: Conceptualization, Formal analysis, Writing (review and editing). **M.Mi.**: Investigation, Software, Writing (review and editing). **N.S.**: Investigation, Software, Writing (review and editing). **Y.S.**: Writing (review and editing). **M.T.**: Writing (review and editing). **K.K.**: Funding acquisition, Writing (review and editing). **K.S.:** Writing (review and editing). **M.E.**: Writing (review and editing). **K.T.**: Writing (review and editing). **H.K.**: Funding acquisition, Writing (review and editing). **S.N.**: Conceptualization, Formal analysis, Funding acquisition, Project administration, Supervision, Writing (original draft), Writing (review and editing). **E.M.**: Conceptualization, Formal analysis, Funding acquisition, Project administration, Supervision, Writing (original draft), Writing (review and editing). All authors contributed to the final manuscript.

## Acknowledgments

The authors would like to thank Sotaro Kikuchi for the useful insights gained in group discussions. The authors also thank Keren-Happuch E Fan Fen for her critical reading of the manuscript.

