## Supplementary Figures and Tables for "Identification of three distinct cell populations for urate excretion in human kidney"

### *Supplementary Material*

#### 1 Supplementary Figures and Tables

##### 1.1 Supplementary Figures

Sakaguchi, Wiriyasermkul, Matsubayashi et al. Supplementary Figure 1

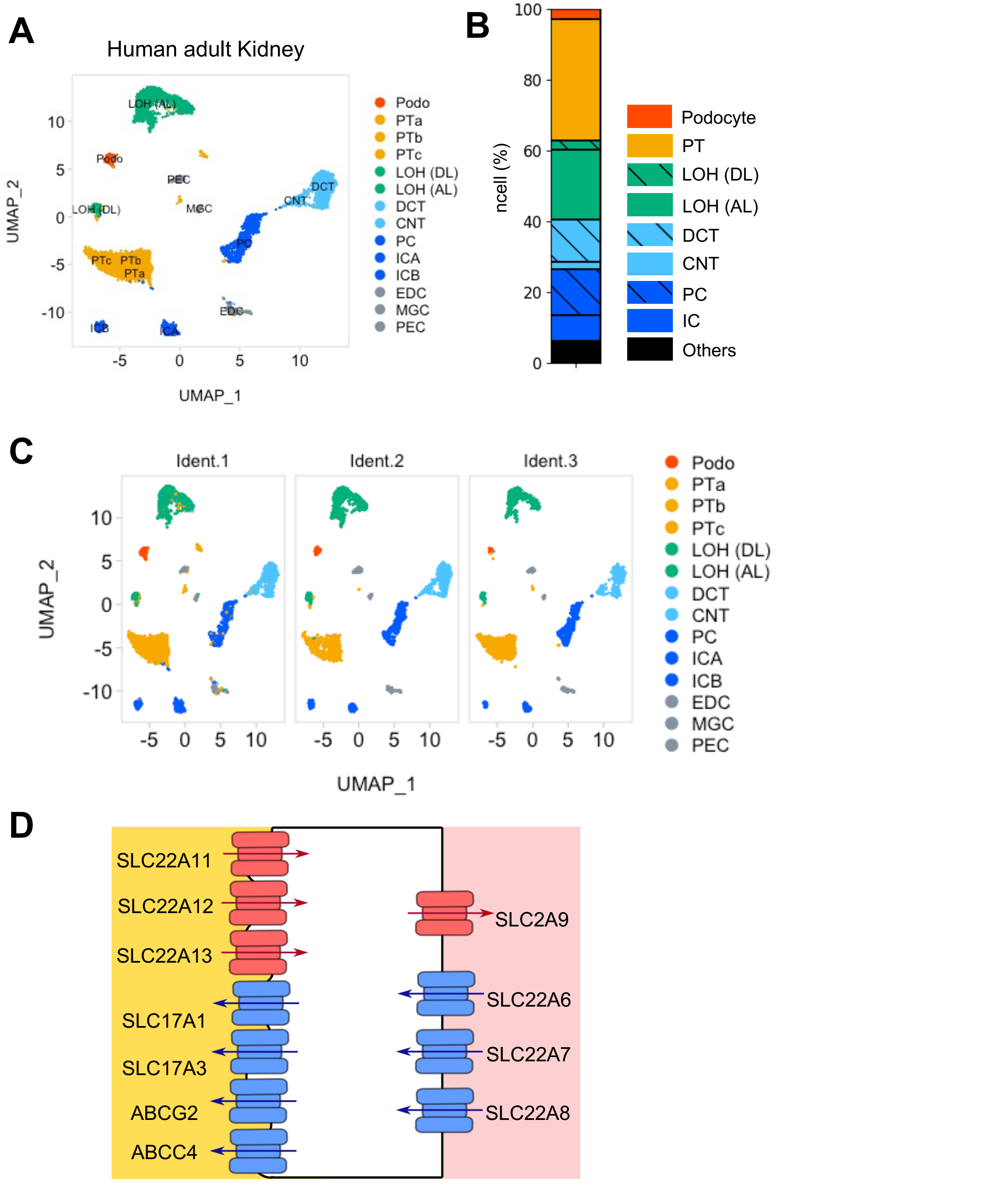

**Supplementary Figure 1. Identification of renal cell types in the datasets.** (A) Unsupervised clustering of healthy human adult renal cells after the annotation. The clusters were named after the anatomical structures of kidney: Podo, podocyte; PT_S1-S3, S1-S3 segments of proximal tubules; LOH (DL), the loop of Henle (descending loop); LOH (AL), the loop of Henle (ascending loop); DCT, distal convoluted tubule; CNT, connecting tubule; PC, principal cell; ICA, intercalated cell type A; ICB, intercalated cell type B; EDC, endothelial cell; MGC, mesangial cell; and PEC, parietal epithelial cell. (B) Proportion of cell clusters of the nephron in all datasets. (C) Unsupervised clustering of three human datasets. (D) A schematic diagram of the subcellular localization of eleven known urate transporters. The apical membrane is to the left of the cells and the basolateral membrane is the right of the cell . Red symbols represent transporters which are reconstituted to urate reabsorption. Blue symbols represent transporters which are reconstituted to urate secretion. Arrows indicate directions of urate flow.

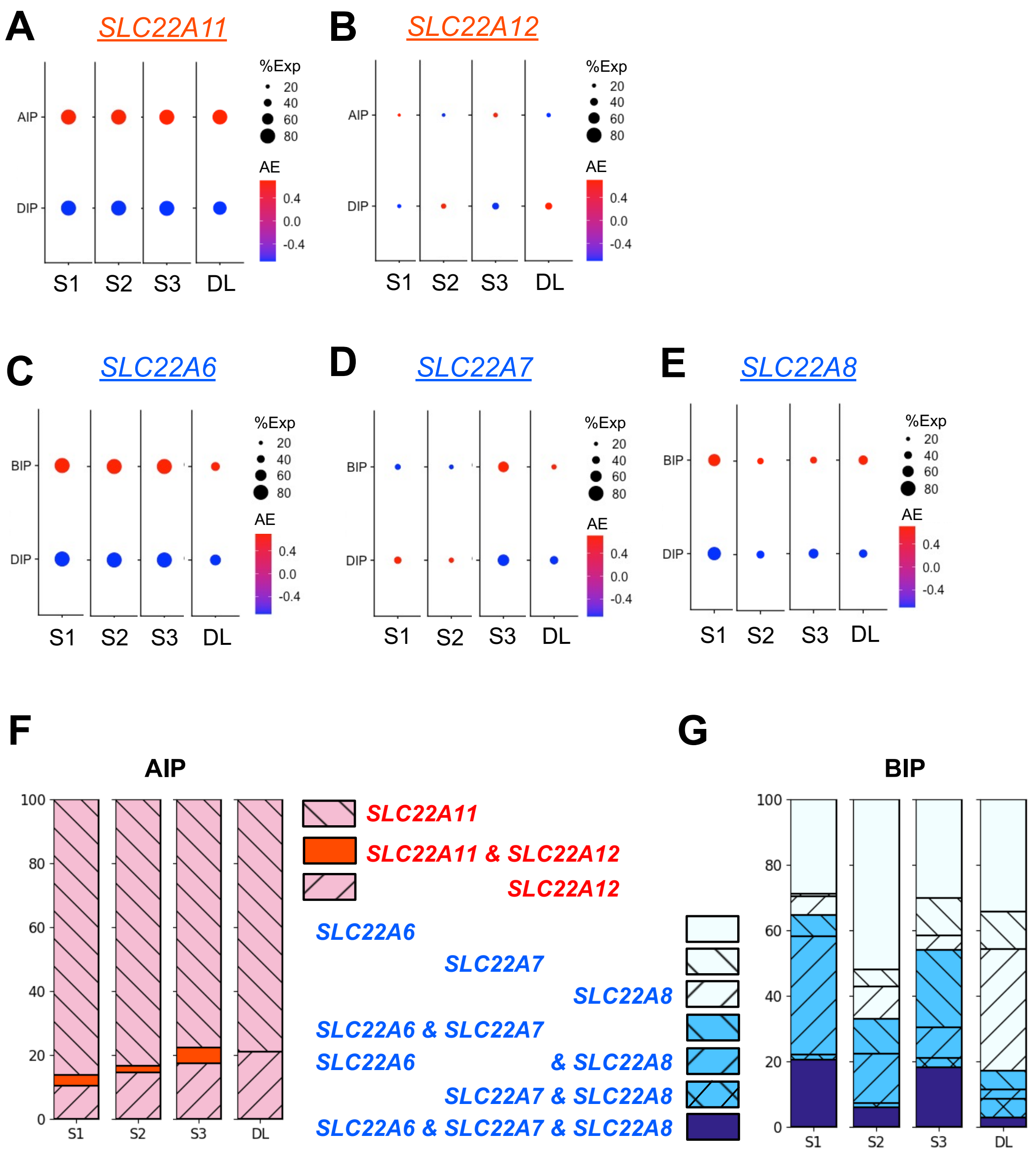


**Supplementary Figure 2. Expression of influx transporters in each cluster.** (A-E) Dot-plots indicate the frequency and the expression levels of urate transporters (x-axis) across the cell populations (y-axis) along the three PT segments (S1-S3) and DL. Dot sizes refer to the frequency of a molecule expressed in the cell population (%Exp), while dot colors indicate the expression levels (AE). SLC22A11(A) and SLC22A12 (B) are the AI transporters (A), described in red letters. SLC22A6 (C), SLC22A7 (D), and SLC22A8 (E) are the BI transporters, described in blue letters. (F-G) Bar plots indicate the percentage (y-axis) of AI transporters in the AIP cell population (F) and BI transporters in the BIP cell population (G) across the regions (x-axis).

Sakaguchi, Wiriyasermkul, Matsubayashi et al. Supplementary Figure 3

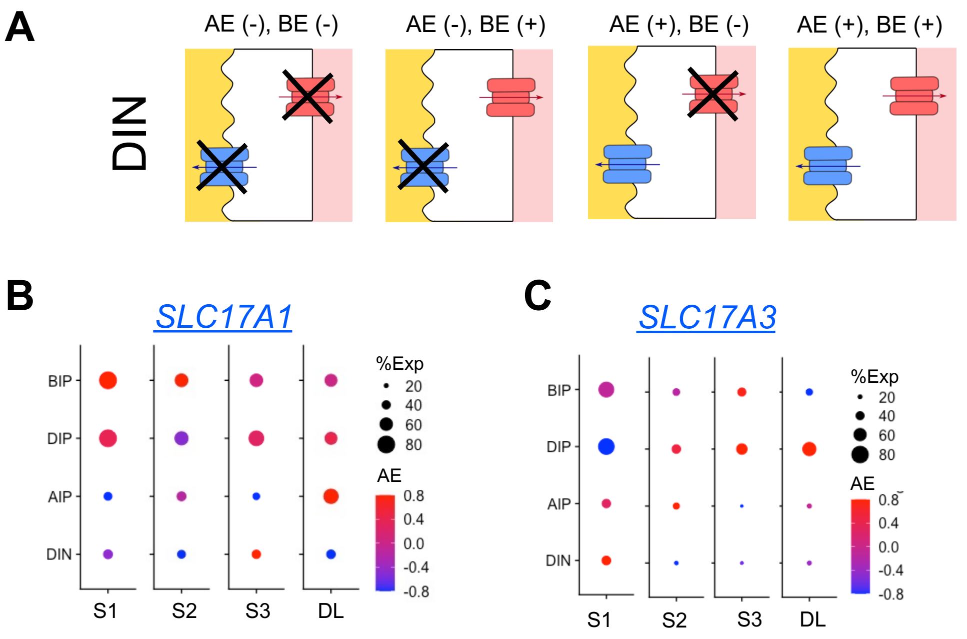


**Supplementary Figure 3. Expression of efflux transporters in each cluster.** (A) Models illustrate the potential expression patterns of urate transporters in the DIN cell population. Colors indicate types of transporters: red, AI or BE transporters; blue, BI or AE transporters. Arrows show urate transport directions. (B-C) Dot-plots indicate the frequency and the expression levels of the AE transporters (B: SLC17A1, C: SLC17A3) (x-axis) across the cell populations (y-axis) along the three PT segments (S1-S3) and DL. Dot sizes refer to the frequency of a molecule expressed in the cell population (%Exp), while dot colors indicate the expression levels (AE).

**A**

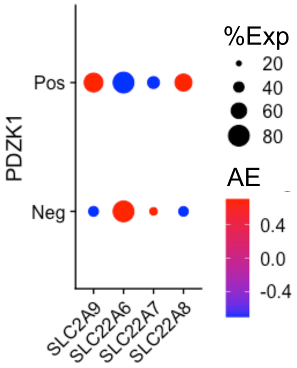


**Supplementary Figure 4. Schematic models of the types of urate transporters expressed in each cell population.** (A) The dot-plot indicates the frequency and the expression levels of the basolateral transporters (x-axis) across the positivity of PDZK1 (y-axis). Dot sizes refer to the frequency of a molecule expressed in the cell population (%Exp), while dot colors indicate the expression levels (AE).

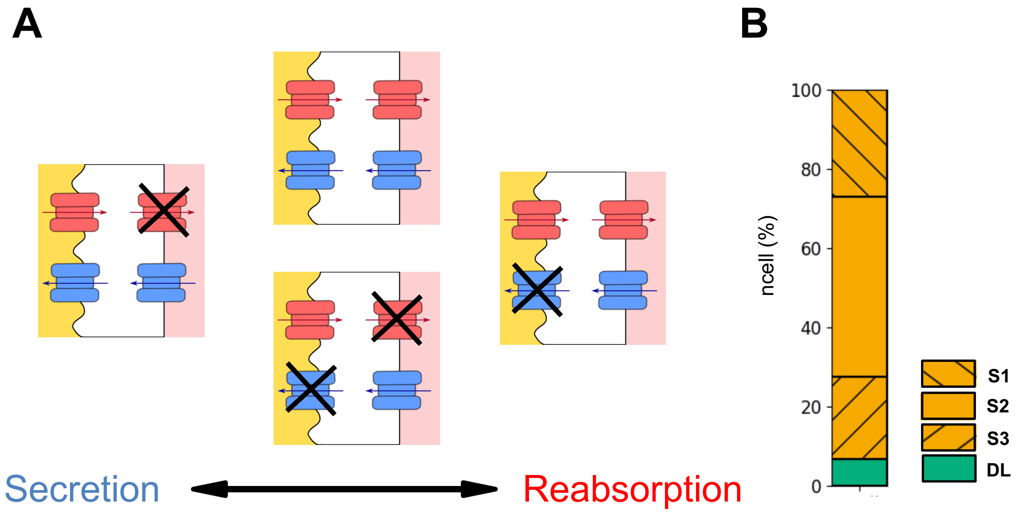


**Supplementary Figure 5. Schematic models of urate transport modes in each cell population.** (A) Representative cell populations contributed to urate reabsorption and secretion in the DIP cell population. Colors indicate types of transporters: red, AI or BE transporters; blue, BI or AE transporters. (B) Proportion of cell clusters in three PT segments and LOH (DL) clusters in the datasets. PT_S1, S1 segment of proximal tubule; PT_S2, S2 segment of proximal tubule; PT_S3, S3 segment of proximal tubule; and LOH (DL), the loop of Henle (descending loop).

#### 1.2 Supplementary Tables

**Supplementary Table 1. Information of snRNA-seq datasets analyzed in this study.**

| Dataset | Species | Age | Sex | GSE | Cell number |
|---|---|---|---|---|---|
| Ident. 1 | Human | 62 years old | Male | GSE118184 | 4524 |
| Ident. 2 | Human | 54 years old | Male | GSE131882 | 3184 |
| Ident. 3 | Human | 62 years old | Male | GSE131882 | 2372 |

**Supplementary Table 2. Marker genes and cell numbers in each renal tubular regional clusters.**

| Cluster name | *Gene* | Cell number |
|---|---|---|
| Podocyte | *NPHS1, NPHS2, PODXL* | 283 |
| PT | *CUBN, LRP2* | 3453 |
| LOH（DL） | *CRYAB* | 257 |
| LOH（AL） | *SLC12A1, UMOD* | 1993 |
| DCT | *SLC12A3* | 1204 |
| CNT | *SLC12A3, CALB1* | 206 |
| PC | *CALB1, ATP1B3, AQP2* | 1304 |
| IC-A | *SLC4A1, AQP6, SLC26A7* | 467 |
| IC-B | *SLC26A4* | 287 |
| EDC | *EMCN, PLAT, ITGA8* | 291 |
| MGC | *ITGA8, PDGFRB* | 75 |
| PEC | *CFH, CLDN1* | 260 |

**Supplementary Table 3. Marker genes and cell numbers in three segments (S1 – S3) of proximal tubule (PT) clusters.**

| Cluster name | *Gene* | Cell number |
|---|---|---|
| S1 | *GLUT2, SGLT2, SLC36A2* | 1000 |
| S2 | - | 1686 |
| S3 | *GLUT1, SGLT1, CaSR, PTH1R* | 767 |

**Supplementary Table 4. Analyses of cell populations in the *Cellular Urate Transport Excretion model*.**

| Region | Population | Type | Counts (n) | Mode | Column "Counts" / Total cell nubmer of each cell population (%) | Column "Counts" / Total cell nubmer (%) |
|---|---|---|---|---|---|---|
| S1 | BIP (46%) | BI | 26 | | 5.9 | 2.6 |
| | | BI +AE | 130 | | 29.4 | 13.0 |
| | | BI +BE | 8 | | 1.8 | 0.8 |
| | | BI +AE+BE | 278 | | 62.9 | 27.8 |
| | DIP (46%) | BI+AI | 25 | | 5.3 | 2.5 |
| | | BI+AI+AE | 126 | | 26.6 | 12.6 |
| | | BI+AI +BE | 11 | | 2.3 | 1.1 |
| | | BI+AI+AE+BE | 311 | | 65.8 | 31.1 |
| | AIP (3%) | AI | 8 | | 27.6 | 0.8 |
| | | AI+AE | 11 | | 37.9 | 1.1 |
| | | AI +BE | 5 | | 17.2 | 0.5 |
| | | AI+AE+BE | 5 | | 17.2 | 0.5 |
| | DIN (5%) | | 16 | | 28.6 | 1.6 |
| | | AE | 31 | | 55.4 | 3.1 |
| | | +BE | 2 | | 3.6 | 0.2 |
| | | AE+BE | 7 | | 1.6 | 0.7 |
| S2 | BIP(43%) | BI | 164 | | 23.4 | 9.7 |
| | | BI +AE | 226 | | 32.2 | 13.4 |
| | | BI +BE | 48 | | 6.8 | 2.8 |
| | | BI +AE+BE | 264 | | 37.6 | 15.7 |
| | DIP (25%) | BI+AI | 67 | | 17.9 | 4.0 |
| | | BI+AI+AE | 149 | | 39.8 | 8.8 |
| | | BI+AI +BE | 29 | | 7.8 | 1.7 |
| | | BI+AI+AE+BE | 129 | | 34.5 | 7.7 |
| | AIP (8%) | AI | 40 | | 39.2 | 2.4 |
| | | AI+AE | 36 | | 35.3 | 2.1 |
| | | AI +BE | 5 | | 4.9 | 0.3 |
| | | AI+AE+BE | 21 | | 20.6 | 1.2 |
| | DIN (24%) | | 225 | | 44.3 | 13.3 |
| | | AE | 166 | | 32.7 | 9.8 |
| | | +BE | 52 | | 10.2 | 3.1 |
| | | AE+BE | 65 | | 12.8 | 3.9 |
| S3 | BIP (41%) | BI | 67 | | 19.8 | 8.7 |
| | | BI +AE | 69 | | 20.4 | 9.0 |
| | | BI +BE | 35 | | 10.4 | 4.6 |
| | | BI +AE+BE | 167 | | 49.4 | 21.8 |
| | DIP (28%) | BI+AI | 37 | | 11.8 | 4.8 |
| | | BI+AI+AE | 63 | | 20.1 | 8.2 |
| | | BI+AI +BE | 27 | | 8.6 | 3.5 |
| | | BI+AI+AE+BE | 186 | | 59.4 | 24.3 |
| | AIP (5%) | AI | 22 | | 55.0 | 2.9 |
| | | AI+AE | 10 | | 25.0 | 1.3 |
| | | AI +BE | 3 | | 7.5 | 0.4 |
| | | AI+AE+BE | 5 | | 12.5 | 0.7 |
| | DIN (26%) | | 32 | | 42.1 | 4.2 |
| | | AE | 24 | | 31.6 | 3.1 |
| | | +BE | 7 | | 9.2 | 0.9 |
| | | AE+BE | 13 | | 17.1 | 1.7 |
| DL | BIP (15%) | BI | 4 | | 11.4 | 1.6 |
| | | BI +AE | 4 | | 11.4 | 1.6 |
| | | BI +BE | 7 | | 20.0 | 2.7 |
| | | BI +AE+BE | 20 | | 57.1 | 7.8 |
| | DIP (6%) | BI+AI | 0 | | 0.0 | 0.0 |
| | | BI+AI+AE | 3 | | 21.4 | 1.2 |
| | | BI+AI +BE | 2 | | 14.3 | 0.8 |
| | | BI+AI+AE+BE | 9 | | 64.3 | 3.5 |
| | AIP (8%) | AI | 4 | | 21.1 | 1.6 |
| | | AI+AE | 8 | | 42.1 | 3.1 |
| | | AI +BE | 1 | | 5.3 | 0.4 |
| | | AI+AE+BE | 6 | | 31.6 | 2.3 |
| | DIN (71%) | | 71 | | 37.6 | 27.6 |
| | | AE | 35 | | 18.5 | 13.6 |
| | | +BE | 27 | | 14.3 | 10.5 |
| | | AE+BE | 56 | | 29.6 | 21.8 |
| | | | | | 1st, 2nd | > 10%, > 5% |
